# Sex-determination cascade orchestrates male-male pheromone synthesis in the bean bug

**DOI:** 10.64898/2026.03.09.710008

**Authors:** Ji-Chong Zhuo, Hai-Qiang Wang, Qing-Ling Hu, Ze-Ping Mao, Lin Wang, Fan Wu, Jia-Bao Lu, Hai-Jian Huang, Zong-Tao Sun, Fei Yan, Jian-Ping Chen, Jun-Min Li, Chuan-Xi Zhang

## Abstract

Volatile pheromones are vital for insect intraspecific communication, yet the genetic basis of homosexual recognition remains elusive. In the bean bug *Riptortus pedestris*, we show that two previously identified aggregation pheromone components, (E)-2-hexenyl-(Z)-3-hexenoate (E2HZ3H) and (E)-2-hexenyl-(E)-2-hexenoate (E2HE2H), serve as key chemosensory cues for male-male recognition during mate selection. Their biosynthesis is governed by the sex-determination cascade *Rpfmd-Rpdsx*. The male-specific isoform *Rpdsx_M* promotes pheromone production in males and induces ectopic synthesis in females upon knockdown of the feminizing switch gene *Rpfmd*. Knockdown of *Rpdsx_M* in males abolishes both compounds, prompting wild-type males to court them as if they were females. Metathoracic gland cells act as the production hub. Behaviourally, E2HZ3H or E2HE2H disrupt mating when applied to females: males avoid such females. E2HZ3H reduces female mobility in the presence of the male-derived primary aggregation pheromone tetradecyl isobutyrate (14:iBu), whereas E2HE2H shows no obvious such effect. These differential effects ensure mating accuracy. The discovery of volatile pheromones functioning in male-male recognition and of their synthesis being governed by the sex-determination cascade updates our understanding of mating accuracy in insect chemical communication.

## Introduction

Volatile sexual pheromones—chemical signals that strongly influence conspecific behavior—represent one of the most widespread and effective strategies for sex recognition and mate location, often exhibiting gender-specific properties that enhance reproductive success^1–5^. In many species, female-produced volatiles primarily guide mate-finding, whereas non-volatile contact pheromones mediate species- and sex-recognition during mate choice^6,7^. Male-produced volatile attractants, commonly classified as aggregation pheromones and composed of low-molecular-weight esters, acids, alcohols, and isoprenoids, can attract both sexes and have been documented across beetles, mites, cockroaches, flies, wasps, bees, thrips, and locusts^8–12^. While a key function of male aggregation pheromones is to attract potential mates, the molecular and genetic mechanisms that enable males to recognize each other during aggregation remain largely unknown.

Evidence from several insect species suggests that homosexual recognition via volatile cues is widespread and functionally diverse. In *Drosophila melanogaster*, the male-produced lipid cis-vaccenyl acetate is transferred to females during copulation, thereby inhibiting sexual advances from other males^13–15^. In *Drosophila grimshawi*, males deposit long-lasting pheromone streaks that attract both sexes to leks, a behavior further enhanced by their ability to detect rival males and increase pheromone deposition to boost female attraction^16,17^. Similarly, in the bed bug (*Cimex lectularius*), alarm pheromones emitted by both sexes serve dual purposes: males use these signals to reduce the risk of homosexual mating, while females employ them to avoid excessive mating attempts^18,19^. Despite these insights, it remains unclear whether males produce volatile pheromones specialized for male–male recognition during mate seeking, and if so, what the underlying molecular mechanisms—particularly the upstream regulators and biosynthetic genes—might be. Filling this gap is essential for a fuller understanding of insect chemical communication.

The bean bug, *Riptortus pedestris* (Hemiptera: Heteroptera: Coreidae), a major pest of soybean fields in East Asia, provides an excellent model for investigating the mechanisms underlying male–male recognition^20,21^. Males produce aggregation pheromones that not only attract females but also males, indicating a sophisticated semiochemical system^22,23^. The primary conspecific attractant, tetradecyl isobutyrate (14:iBu), was reported to exhibit enhanced behavioral efficacy when combined with synergistic compounds—(E)-2-hexenyl hexanoate (E2HH), (E)-2-hexenyl (Z)-3-hexenoate (E2HZ3H), (E)-2-hexenyl (E)-2-hexenoate (E2HE2H), and octadecyl isobutyrate^24,25^. However, E2HZ3H and E2HE2H showed no attractiveness when applied individually, and their effect did not correlate with the doses added to 14:iBu^26^. Moreover, among the attracted individuals, most males lack 14:iBu but instead contain E2HZ3H and E2HE2H^27^. Since the presence of E2HZ3H and E2HE2H is restricted to sexually mature males, these compounds likely play a specific role in sexual communication.

Here, we show that E2HE2H and E2HZ3H function as male–male recognition pheromones and, by regulating mating behavior, help ensure accurate mate recognition. Mechanistically, we demonstrate that the sexual dimorphism regulator *doublesex*(*dsx*) controls mating behavior by directing the biosynthesis of these two pheromones. Importantly, both compounds hold substantial potential as behavior-based mating disruption agents for environmentally friendly pest control strategies.

## Methods

### Insect Husbandry

The bean bug (*Riptortus pedestris*) strain used in this study was originally collected in a soybean field in Suzhou, China in 2019. The insects were reared in separate chambers(30 x 30 x 30 cm) with soybean plants (strain: Wandou 27) at 26 ± 0.5°C, with a 50 ± 5% relative humidity and a 16:8 h light/dark photoperiod. For harvesting pools of synchronized bean bug, freshly emerged adults were separated by sex on the day of moulting (day 0) and placed female and male adults in different 500 mL plastic cups. They were provided with soybeans and tap water.

### Gene cloning

Gene-specific primers were designed from the transcriptome sequences to amplify the full open reading frame (ORF) of *Rpdsx_F*, *Rpdsx_M, Rpfmd*. The PCR products were cloned into a PMD-19T vector (TaKaRa, Dalian, China) and sequenced. The sequence-verified strains were preserved at -80°C.

### RNA interference

The gene-specific coding sequences of *Rpdsx*, *Rpfmd*and green fluorescent protein (*gfp*, as a control) were cloned into PMD-19T vectors. PCR-generated DNA templates were subsequently used to synthesize dsRNA containing T7 promoter sequences at both ends. Gene-specific dsRNAs were synthesized using a MEGAscript T7 transcription kit (Ambion, Austin, TX, USA) and quantified spectrophotometrically with a NanoDrop 2000 (Thermo Fisher Scientific, Franklin Lakes, NJ, USA). The quality and size of the dsRNA were verified by 1% agarose gel electrophoresis. The final dsRNA concentration was approximately 3–5 μg/μL. Microinjections were performed by administering 0.2 μL of dsRNA into third-instar nymphs and 2 μL into female adults. The dsRNA injection procedures for *R. pedestris* followed methods previously described for brown planthopper (BPH) dsRNA treatments by Xu et al (https://doi.org/10.1038/protex.2015.005).

### Transcriptomic sequencing and analysis

For second generation Illumina sequencing (3 biological, each with 5 technical replicates), the total RNAs of female and male adults were extracted with Trizol. The total RNA was delivered to the Novogene Company (Tianjin, China) for Illumina 150 bp paired-end sequencing on a HiSeq4000 platform. The clean reads were aligned to the *R. pedestris* reference genome using HISAT2 v2.1.0^21^. The count data of each gene based on the genome annotation file was obtained using CUFFQUANT. The differentially expressed genes (DEGs) were identified using DESeq2. Gene expression level was presented as Transcripts per million (TPM) which was calculated using RSEM.

### Single-cell RNA sequencing and analysis

For single-cell RNA sequencing, the metathoracic glands were dissected from male adults and delivered to the Oebiotech Company (Shanghai, China). The protocols for analyzing single-cell RNA sequencing data were used as previously reported^28^. Briefly, The reads were aligned to the *R. pedestris* reference transcriptome using kallisto v0.43.1. The alignment file was quantified using bustools v0.45.1. The count data was loaded into R v4.4.2 using BUSpaRse v1.20.0. The empty droplets were filtered using DropletUtils v 1.26.0. The cell type atlas of the metathoracic gland was created using Seurat v5.3.0 and Harmony v 1.2.3.The gene expression programs (GEPs) were inferred using consensus Non-negative Matrix Factorization (cNMF) in Python v 3.9.21.

### RNA extraction, qRT-PCR and semiquantitative RT-PCR

Total RNAs were extracted from the various tissues and different developmental stages using the TRIzol reagent (TaKaRa) according to the manufacturer’s recommendations and immediately stored at -80 °C. The quantity of the total RNA was assessed using a NanoDrop One spectrophotometer (Thermo Fisher Scientific). 1μg of the total RNA was reverse-transcribed to first-strand cDNA using the HiScript® II Q RT SuperMix for qPCR (+gDNA wiper) (Vazyme, Nanjing, China). All qPCR analyses were performed using Hieff® qPCR SYBR Green Master Mix Low Rox Plus (Yeasen, Shanghai, China) with biological replicates. The actin gene was used as an endogenous control to normalize the expression levels. The *Rpdsx* sex-specific transcripts in male and female adults were examined by semiquantitative RT-PCR.

### Metathoracic gland volatile collection and quantification

The volatiles of the virgin males and females were collected and analyzed by headspace solid-phase microextraction (HS-SPME) coupled with gas chromatography-mass spectrometry (GC−MS)^29^. Three *R. pedestris* adults with same stages were placed in 20 mL borosilicate glass volatile organic analysis vials and sealed using Parafilm and placed in -80 °C for 10 min. The various tissue types were dissected from male or female adults (9 d postemergence) and the freeze step was omitted. Subsequently, the sample was dipped in a water bath maintained at 25 °C to thaw for 5 min. Finally, the SPME extraction was performed for 1 h. Purple SPME fibers (polyethylene glycol, 60 μm) and manual holder (Supelco) were used to collect the volatiles. Before the first time of use, the fibres were immersed in a solution of 1% methanol and 15% sodium chloride for 15–30 min, and then thermally cleaned in the GC injection port at 240 °C for 30 min.

An Agilent 8890A gas chromatograph with a HP-5MS UI capillary column (30 m x 250 μm i.d., 0.25 μm film thickness) and an Agilent 5977B mass selective detector was used to analyse the volatile compounds on the SPME fiber. The inlet temperature operated in splitless mode was maintained at 280 °C with helium as the carrier gas at a constant flow rate of 1.2 mL/min. The fibre was inserted into the inlet and held for 5 min. The GC temperature was programmed at an initial temperature of 50 °C and held for 2 min, increased to 190 °C at a rate of 8 °C/min and held for 1 min, and then increased to 280 °C at a rate of 20 °C/min and held for 5 min. The mass spectrometer was operated in the electron ionization (EI) mode at 70 eV with ion source at 230 °C, with a scan rate of 1.4 scans/sec from m/z 50 to 600. Compounds were tentatively identified by comparisons of their spectral data with mass spectral data from the National Institute of Standards and Technology (NIST23) library, and further confirmed by comparisons of retention times and mass spectra with those of standards. The authentic compounds purchased from chemical companies were used to develop the four-point calibration curves for external standardization and quantification of the volatiles.

### EAG analysis

For EAG measurement, one antenna was cut off from the base of the scape. Then, the tip of the distiflagell was finely excised. Subsequently, the antenna was connected to a glass electrode. This assembly was then connected to a conductive silver wire using a 0.9% sodium chloride solution. A constant airflow (1.5L/min) from a stimulating air stream generator (DGST-2, Syntech, Buchenbach, Germany) was filtered through activated carbon. The filtered stimulation airflow (0.5 L/min) was passed over filter paper containing 10 μL of authentic compound solutions at different concentrations for a duration of 0.5s. To avoid sensory adaptation, two stimulation airflows were separated by 1 min intervals. The EAG signals were amplified by DG3FA-1 (Syntech, Buchenbach, Germany) and were recorded by Keysight BenchVue software (Syntech, Buchenbach, Germany). To check the response to E2HH, E2HZ3H and E2HE2H, ninety antennae from ninety *R. pedestris* were tested.

### Behavioral assays

Single-pair courtship trials were conducted in a plastic dish containing one male and one female *R. pedestris*. The copulation latency was defined as the time from the introduction of the male into the dish to the ovipositor extrusion. Single-pair courtship (male–female) trials were conducted in a 9 cm diameter × 1.5 cm dish containing one 9-day-old wildtype male and one treated female *R. pedestris*. Each experimental treatment consisted of 10 male individuals were tested with an equal number of the treated females. Single-pair courtship (male–male) trials were conducted in a 3.5 cm diameter × 1.5 cm dish containing one 9-day-old wildtype male and one treated male *R. pedestris*. The mating behavior were recorded by mobile phones.

Experiment on *R. pedestris* avoidance by E2HH, E2HZ3H and E2HE2H, one wildtype female or male was put in one 9 cm diameter × 1.5 cm dish, in whose center 0.5cmX0.5cm filter paper was put with 10μL 10% E2HH, E2HZ3H or E2HE2H. The behavior of *R. pedestris* was recorded by a mobile phone and analyzed by sofrware ToxTrac.

## Results

### Male-specific pheromones play critical roles in the mating behavior of *R. pedestris*

Adult male *R. pedestris* produce two sex-specific volatiles, E2HZ3H and E2HE2H, alongside the non–sex-specific compound E2HH. We compared their emission profiles to assess timing and abundance. Volatiles emitted by 7–9-day-old virgin males and females were analyzed using headspace solid-phase microextraction coupled with gas chromatography–mass spectrometry (HS-SPME–GC/MS) (Fig. 1a; Extended Data Fig. 1a, b). Absolute quantities of each compound, which were present at submicrogram levels, were determined using an external standard calibration method (Extended Data Fig. 1c). Quantitative analyses showed that E2HH was produced at comparable levels in males and females, with no significant intersexual difference. In contrast, E2HZ3H and E2HE2H are the male-specific compounds; the former was present in substantially higher amounts than the latter (Fig. 1b). To examine the developmental timing of volatile production, body extracts from 5th-instar nymphs and adults at different ages were analyzed. None of the three compounds were detected in either male or female 5th-instar nymphs (Extended Data Fig.2). E2HH production began on the first day after adult emergence, increased progressively in both sexes, and remained at high levels from day 6 to day 12. By contrast, E2HZ3H and E2HE2H first became detectable on day 3 of male adulthood and reached stable levels between day 6 and day 12, coinciding with the period when mating behavior is typically observed (Fig. 1c,d).

**Figure 1.**
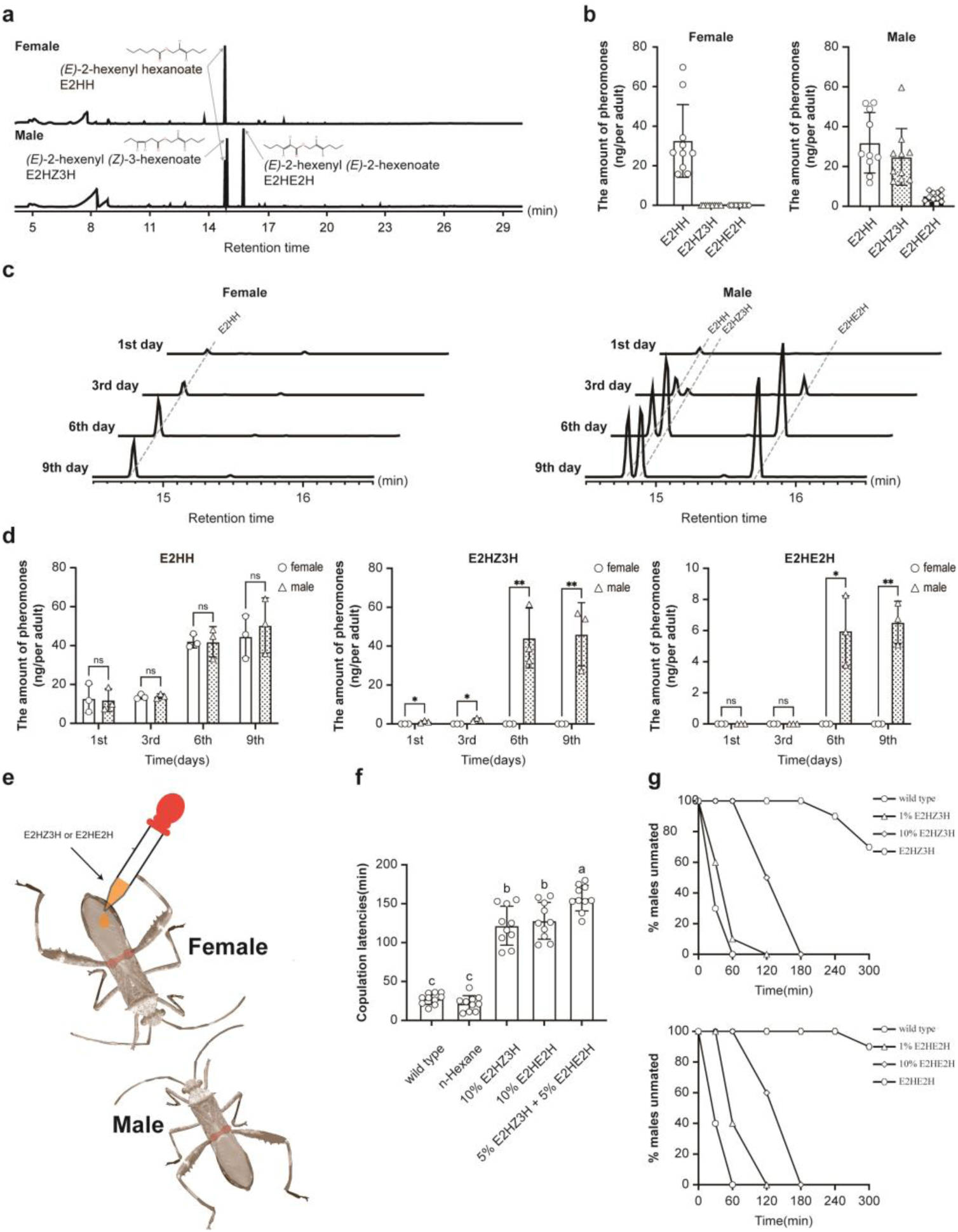
Sex pheromones of *R. pedestris*. (a) Volatile compound profiles of females and males are shown. HS-SPME–GC–MS analysis detected only E2HH in 7-day-old females, whereas E2HH, E2HZ3H, and E2HE2H were identified in 7-day-old males. (b) Quantification of E2HH, E2HZ3H, and E2HE2H in females and males is presented. E2HZ3H is more abundant than E2HE2H in males. Each measurement included more than 10 biological replicates. (c) Temporal dynamics of E2HH, E2HZ3H, and E2HE2H in adult females and males are examined. HS-SPME–GC–MS analysis was performed on females and males at days 1, 3, 6, and 9 after adult emergence. (d) Quantification of E2HH, E2HZ3H, and E2HE2H using the external standard method is shown. Three biological replicates were performed per time point. (e) A schematic diagram illustrates the mating assay. (f) The effects of E2HZ3H and E2HE2H on mating behavior are evaluated in 7-day-old virgin females and males, with more than 10 biological replicates per treatment. (g) Copulation latency of wild-type males paired with pheromone-treated females is quantified. The effects of different concentrations of E2HZ3H and E2HE2H on mating success were assessed over a 300-min observation period, with 10 mating pairs per treatment. Statistical significance was assessed using Student’s t-test (ns, p > 0.05; ***, p < 0.001) and Tukey’s post hoc test (same lowercase letters indicate no significant difference, p > 0.05; different letters indicate significant differences, p < 0.05).

**Figure 2.**
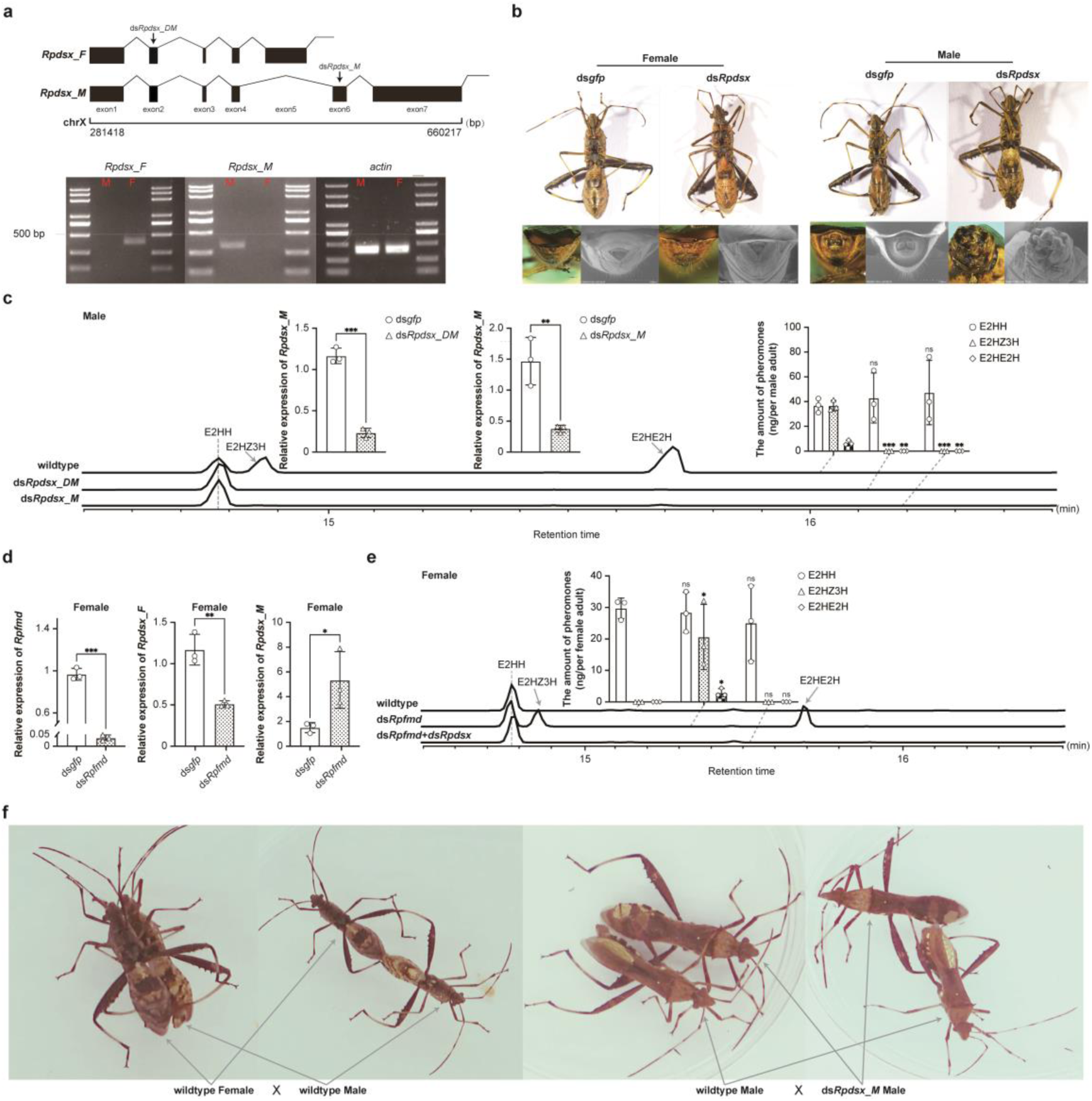
Regulation of the sex-determination cascade on sex pheromone production in *R. pedestris*. (a) Alternative splicing and sex-specific expression of Rpdsx are depicted. The Rpdsx gene produces two sex-specific isoforms, *Rpdsx_M* (exons 1–4, 6, and 7) and *Rpdsx_F* (exons 1–5), and is located on chromosome X (281,418–660,217 bp). Semi-quantitative PCR (35 cycles) reveals female- and male-specific expression patterns. (b) The effects of *Rpdsx* knockdown on somatic development are examined. RNAi targeting the common regions of *Rpdsx_M* and *Rpdsx_F* (ds*Rpdsx_DM*) in third-instar nymphs does not affect female somatic development but causes malformation of male external genitalia. (c) The role of *Rpdsx* in sex pheromone production is assessed. Two independent dsRNAs efficiently knock down *Rpdsx_M* expression in males (>70% efficiency). Both E2HZ3H and E2HE2H are abolished in ds*Rpdsx*-treated males, whereas E2HH levels remain unchanged. (d) Regulation of *Rpdsx* expression and pheromone biosynthesis by *Rpfmd* is demonstrated. ds*Rpfmd* achieves >95% knockdown efficiency, and RNAi of *Rpfmd* in females induces ectopic expression of *Rpdsx_M*. (e) Pheromone biosynthesis controlled by *Rpfmd* is illustrated. E2HZ3H and E2HE2H are detected in ds*Rpfmd*-treated females but become undetectable after subsequent ds*Rpdsx* injection. (f) Representative images illustrate mating behavior. Wild-type females mate with wild-type males, and wild-type males also exhibit mating behavior toward ds*Rpdsx*-treated males. All quantitative analyses were performed with three biological replicates. Statistical significance was assessed using Student’s t-test (ns, p > 0.05; ***, p < 0.001).

To elucidate the functional roles of male-specific compounds (E2HZ3H and E2HE2H) in *R. pedestris* mating behavior, we conducted behavioral assays using 7-day-old virgin females treated with five experimental conditions (1μL/individual): wildtype control, 100% n-hexane solvent control, 10% (v/v) E2HZ3H/n-hexane, 10% (v/v) E2HE2H solution, and 5% E2HZ3H+5% E2HE2H binary blend (Fig. 1e). Standardized mating trials under controlled conditions (25±1°C, 60% RH) revealed that control pairs achieved copulation within 25 minutes via stereotypic behavioral sequences including male mounting, antennal tapping with foreleg vibrations, and successful genital insertion (Supplementary Video. 1). Both E2HZ3H and E2HE2H independently applied onto females induced complete mating failure within 60-minute observations by disrupting sexual recognition, with males exhibiting no mating behavior, and normal recognition and copulation only resumed after the volatile E2HZ3H and/or E2HE2H applied onto females had fully evaporated (Supplementary Video. 1). Quantitative analysis indicated concentration-dependent effects, with individual compounds prolonging mean copulation latency to about 120 minutes (3-fold increase versus controls), while the binary blend showed synergistic disruption extending latency to 150 minutes (Fig. 1f; Supplementary Video. 1). These effects were strongly concentration dependent, with increasing doses leading to a progressively stronger inhibition, including a marked suppression of male mating success (Fig.1g). These results demonstrate that the male-specific volatiles E2HZ3H and E2HE2H are critical for intersexual recognition in *R. pedestris* mating systems.

### The *Rpfmd-Rpdsx* cascade regulates sexual pheromones biosynthesis in adult stage

The gene *doublesex* (*dsx*) plays a pivotal role in sex-specific development in insects, prompting us to investigate whether *dsx* of *R. pedestris* (*Rpdsx*) contributes to the production of sex-specific pheromones and influences mating behavior. In this study, *Rpdsx* was first identified on the X chromosome and found to produce sex-specific isoforms—female-specific *Rpdsx_F* and male-specific *Rpdsx_M*—through alternative splicing (Fig. 2a). RNAi knockdown of *Rpdsx* in 3rd-instar nymphs targeting the common exons of *Rpdsx_F* and *Rpdsx_M* (ds*Rpdsx_DM*) resulted in malformed male external genitalia, with no apparent effect on female somatic development (Fig. 2b). Importantly, the male-specific pheromones E2HZ3H and E2HE2H were absent in emerged males following *Rpdsx* knockdown, whereas the non-sex-specific pheromone E2HH remained unchanged; similarly, RNAi targeting the male-specific exons of *Rpdsx_M* (ds*Rpdsx_M*) produced identical results, indicating that *Rpdsx* not only regulates sex-specific somatic development in the nymph stage but also controls the synthesis of male-specific pheromones in adults (Fig. 2c; Extended Data Fig. 3-4).

In *R. pedestris*, the feminizing switch gene homolog *Rpfmd*, via its female-specific isoform (*Rpfmd_F*), plays a key role in sex differentiation^30^. In the current research, RNAi knockdown of *Rpfmd-F* in females reversed the sex-specific expression of *Rpdsx*, leading to the expression of the male-specific *Rpdsx_M* while reducing the expression of female-specific *Rpdsx_F* (Fig. 2d). Consequently, male-specific pheromones E2HZ3H and E2HE2H were detectable in these females, likely due to *Rpdsx_M* expression, and disappeared again when *Rpdsx_M* was subsequently knocked down via RNAi (Fig. 2e; Extended Data Fig. 4). Collectively, these findings demonstrate that *Rpdsx* regulates the synthesis of E2HZ3H and E2HE2H through its male-specific isoform, and that this regulation is controlled by the upstream feminizing switch *Rpfmd*, forming a *Rpfmd*–*Rpdsx* cascade that governs the male-male recognization pheromone biosynthesis in adults.

Based on the aforementioned research, we observed that ds*Rpdsx_M* males exhibit a volatile compound profile similar to that of wildtype females, raising the question of whether males could recognize them accurately. When 1st-day virgin males were treated with ds*Rpdsx_M* which likewise abolished the production of E2HZ3H and E2HE2H, and then paired with wild-type males, the latter displayed mating behaviors—including mounting, antennal tapping, and body shaking—toward ds*Rpdsx_M* males, indicating misrecognition (Fig. 2f; Supplementary Video.2). To further investigate, we applied E2HZ3H or E2HE2H to the treated males, which enabled wildtype males to make correct identifications and refrain from exhibiting mating behavior. This phenomenon may persist up to 24 hours, until the E2HZ3H or E2HE2H had fully dissipated, wildtype males resumed displaying mating behavior toward the ds*Rpdsx_M* males (Supplementary Video.2).

These findings indicate that both E2HZ3H and E2HE2H serve as male-male recognization pheromones, enabling males to identify individuals of the same gender, and that the sex determination cascade regulates mating behavior by controlling the synthesis of these male-specific pheromones through the male-specific alternative splicing of *Rpdsx*.

### Male-male recognization pheromones are produced in the male metathoracic scent glands and detected by the antennae

Although *R. pedestris* males possess a well-developed metathoracic gland, and such glands in other heteropteran bugs can produce volatile secretions, there is currently no published evidence identifying the glandular source of E2HZ3H and E2HE2H(Fig. 3a). To explore the secretory organization of these pheromones, we performed detailed analyses on the head, thorax, and abdomen of 9-day-old males. Our initial findings indicated that three volatile compounds were exclusively detected in the thorax, where the metathoracic glands are notably prominent(Fig. 3b). To further ascertain whether the specific compounds E2HH, E2HZ3H, and E2HE2H were synthesized within these glands, we conducted separate analyses of the metathoracic glands and the thorax with the glands excised. The results revealed that the majority of these compounds were predominantly concentrated in the metathoracic glands, providing strong evidence that they serve as the primary synthesis sites for E2HH, E2HZ3H, and E2HE2H (Fig. 3b). To gain deeper insights into the cellular mechanisms underlying pheromone production, we employed single-cell RNA sequencing (scRNA-seq) using droplet microfluidics to comprehensively profile the entire metathoracic gland (Fig. 3c). A total of 12 distinct cell types within the gland were identified (Fig. 3d). The co-expression pattern of key related genes strongly suggests that cell type 9 functions as a specialized secretory cell type (Extended Data Fig.5). Collectively, these findings indicate that the male-male recognization pheromones are biosynthesized in the metathoracic gland.

**Figure 3.**
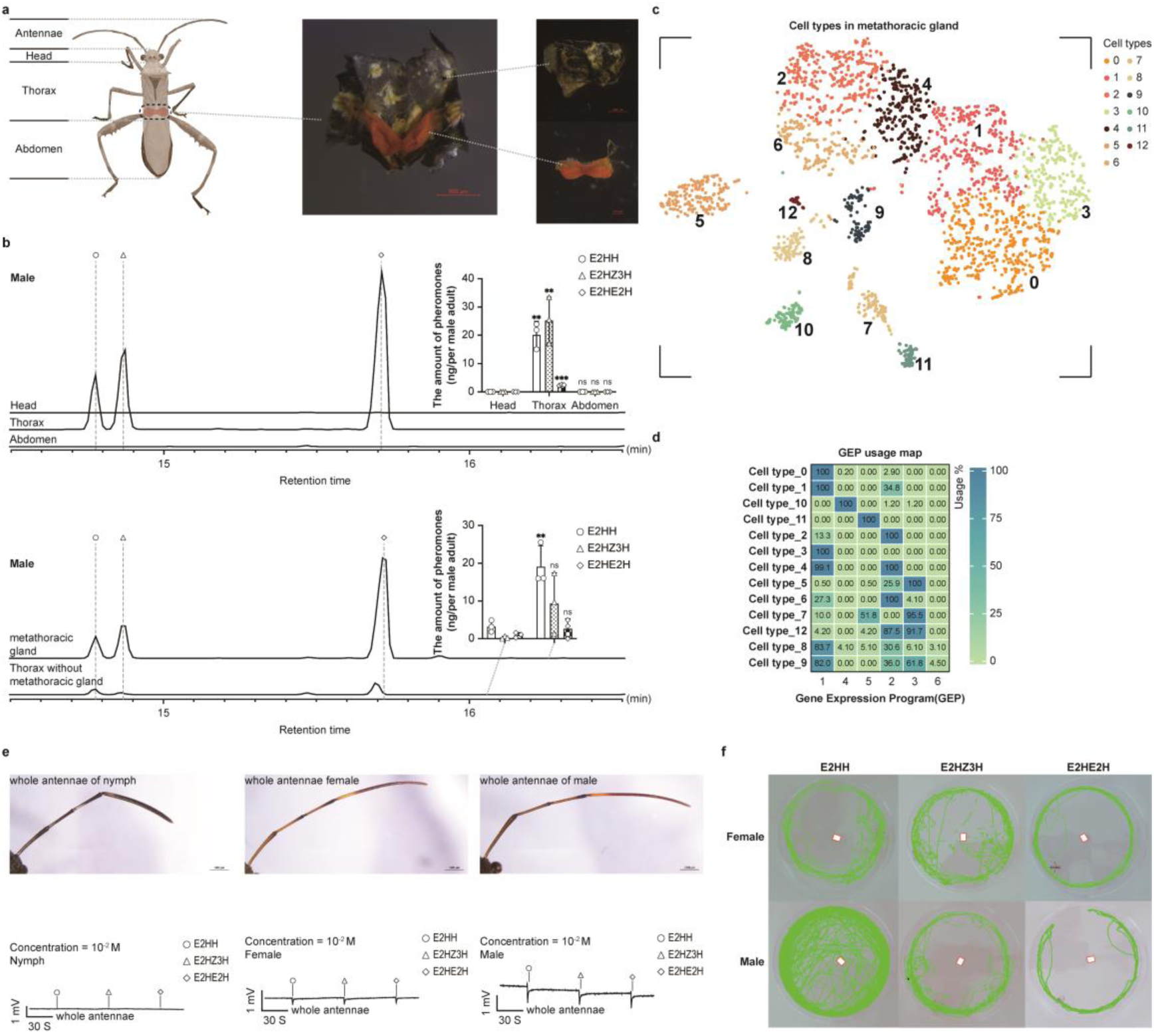
Synthesis and detection of sex pheromones in *R. pedestris*. (a) The biosynthesis of E2HH, E2HZ3H, and E2HE2H occurs in the metathoracic gland. Dissected head, thorax, and abdomen are shown, with the metathoracic gland highlighted in orange together with other thoracic tissues. (b) E2HH, E2HZ3H, and E2HE2H are predominantly detected in the thorax, and their quantities were measured. The levels of these pheromones are significantly higher in the metathoracic gland than in other thoracic tissues. (c) A single-cell atlas of the metathoracic gland is shown. UMAP visualization reveals distinct cell-type clusters. (d) Gene expression program (GEP) usage across 12 cell categories is shown. (e) Detection of pheromones across developmental stages and sexes is shown using antennae from nymphs, females, and males. Electroantennography (EAG) recordings show antennal responses to E2HH, E2HZ3H, and E2HE2H at 10⁻² M. (f) Behavioral responses of wild-type females and males to pheromones are shown. Individuals were tested in 9 × 1.5 cm arenas, and movement trajectories were recorded for approximately 180 min. Filter papers (0.5 × 0.5 cm) containing 10 μL of 10% E2HH, E2HZ3H, or E2HE2H were placed at the center of the arenas (red rectangles), and green lines indicate movement tracks. Statistical significance was determined using Student’s t-test (ns, p > 0.05; **, p < 0.01; ***, p < 0.001).

Given that mating in *R. pedestris* ony occurs between sexually mature females and males, and that male-specific sex pheromones are synthesized during the adult stage. We hypothesized that these compounds would display both sex- and stage-specific perceptual characteristics. Previous studies have shown that the antennae are the principal sensory organs in *R. pedestris* for detecting such chemical signals. To test this, we conducted electroantennography (EAG) to evaluate the antennal responses of males, females, and fifth-instar nymphs to E2HH, E2HZ3H, and E2HE2H (Fig. 3e; Extended Data Fig.6). Our results demonstrated that both adult males and females exhibited clear antennal responses to all three compounds, indicating their sensitivity to these pheromones. By contrast, the antennae of fifth-instar nymphs showed no significant responses, suggesting that E2HZ3H and E2HE2H function specifically in chemical communication between adults. In behavioral assays, females showed avoidance toward filter papers treated individually with E2HH, which also produced by females, a response that may contribute to population dispersal (Fig. 3f). Females also avoided the male-specific E2HZ3H, or E2HE2H respectively, and we found that the existence of 14:iBu could offset the corresponding avoidance effect, moreover, 14:iBu and E2HZ3H could significantly decreases the mobility of female insects, which is consistent with the natural behavior of females staying in place to wait males for mating,whereas E2HE2H shows no obvious such effect. (Supplementary Video1; Fig. 3f; Extended Data Fig. 7). Considering that Males also avoided filter papers treated with E2HZ3H and E2HE2H, no matter whether 14:iBu existed or not, supporting their function as male–male recognition pheromones. In contrast, E2HH did not induce avoidance in males (Fig. 3f, Extended Data Fig.7). Considering that E2HH is also produced by females, its mere presence may function as a female-specific signal. Together, these findings suggest that the selective perception of such compounds monimizes nymphal interference, thereby enhancing both mate recognition accuracy and reproductive isolation among adults.

## Discussion

Insect sexual communication relies heavily on chemical codes that allow individuals to locate and identify suitable mates. Although many species use long-range attractants or contact hydrocarbons for this purpose, it remains unknown whether volatile pheromones specialized for male–male recognition exist, and if so, how they are regulated at the molecular level. In *R. pedestris*, male-derived aggregation pheromones attract both wild males and females, potentially compromising precise mate recognition^22^. Our work identifies two male-specific volatiles—E2HZ3H and E2HE2H—as the core constituents of the recognition code in this species. These volatiles are both necessary and sufficient for males to distinguish females from males: applying them to virgin females disrupts this discrimination, whereas their absence in ds*Rpdsx* males leads wild-type males to misidentify these males as females, a phenotype fully restored upon reintroducing either compound. Together, these findings uncover a dedicated volatile module that chemically encode sexual identity, ensuring accurate mate recognition even in complex social and ecological settings.

Male-produced sex pheromones are widely documented across multiple insect orders, including Coleoptera, Diptera, Hemiptera, Lepidoptera, and Dictyoptera, where they serve as critical sex attractants, yet the molecular mechanisms controlling their male-specific expression remain poorly understood, especially for pheromones involved in male–male recognition^9,31–35^. In this study, we uncover volatile pheromones specific serve as key chemsosensory cues for male-male recognition during mate selection and a previously uncharacterized regulatory mechanism whereby the sex-determination cascade, specifically the *Rpfmd*–*Rpdsx* axis, governs the biosynthesis of male-specific volatile pheromones that mediate male–male recognition in *R. pedestris*. We show that the male-specific isoform *Rpdsx_M* acts as a positive regulator of E2HZ3H and E2HE2H, which are secreted exclusively by males. RNAi knockdown of *Rpfmd* induces ectopic *Rpdsx_M* expression in females, leading to de novo production of these male-specific pheromones, whereas subsequent silencing of *Rpdsx_M* abolishes this induced synthesis, demonstrating that *Rpdsx_M* is both necessary and sufficient for activating the biosynthesis of male-emitted pheromones involved in male–male recognition. This regulatory logic contrasts with other insects: in *Blattella germanica*, male-specific *dsx* isoforms suppress pheromone biosynthesis^28^; in *Bactrocera dorsalis*, male *dsx* represses elongase11 while female *dsx* promotes pheromone production^36^. Thus, our findings reveal an innovative, lineage-specific mode of *dsx*-mediated control, in which a male *dsx* isoform actively drives the production of volatile male-specific pheromones used for male–male signaling. This work provides new insight into the evolution of insect chemical communication and expands the functional diversity known for the *dsx* gene.

In summary, we show that the male-specific isoform *Rpdsx_M* directs biosynthesis of E2HZ3H and E2HE2H, pheromones that define male identity and mediate male–male recognition. This work reveals a lineage-specific, active role of doublesex in shaping male chemical communication, linking molecular regulation, synthesis hubs, and behavioral output (Fig. 4). By connecting sex-determination cascades to pheromone evolution, we provide a conceptual framework for understanding the diversification of sex-specific chemical signaling in insects, highlighting how genetic control generates species-specific cues. Our findings expand the known functions of volatile pheromones beyond attraction and mate location, establishing a novel role in male–male recognition. Moreover, we present the first example of an insect sex-determination cascade directly regulating volatile homosexual pheromone production, integrating developmental identity with adult sexual interaction.

**Figure 4.**
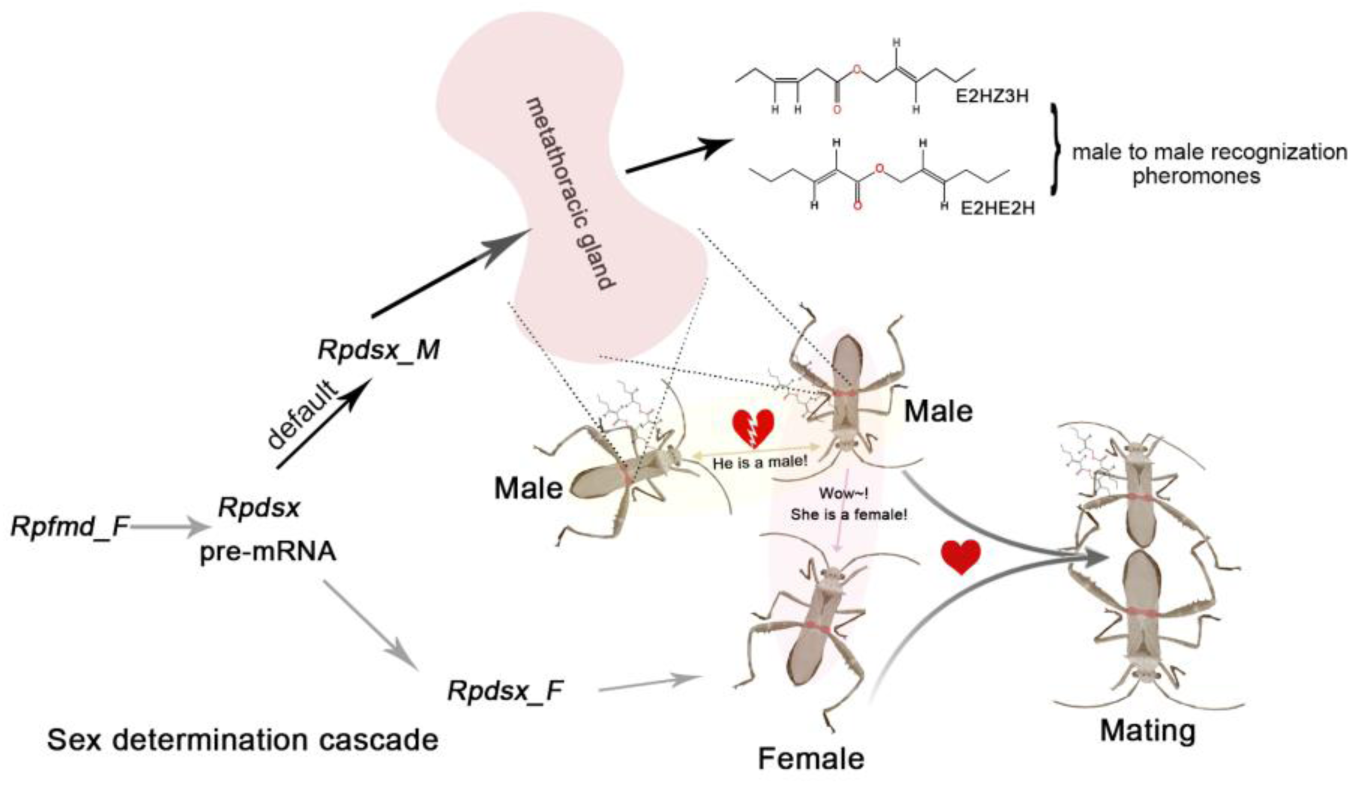
The sex determination cascade governs the biosynthesis of male–male recognition pheromones in *R. pedestris*. In males, the male-specific isoform *Rpdsx_M* is produced by default via alternative splicing of *Rpdsx* pre-mRNA. RpDSX_M promotes the biosynthesis of E2HZ3H and E2HE2H in the metathoracic gland. In females, *Rpfmd_F* controls the alternative splicing of *Rpdsx* pre-mRNA to generate the female-specific isoform *Rpdsx_F*. During mating, males rely on E2HZ3H and E2HE2H to distinguish females from other males, enabling them to initiate courtship and achieve successful copulation. E2HZ3H and E2HE2H together provide a double-check mechanism to ensure mating accuracy.

## Data availability

All data that support the findings of this study are available within the Article and its Supplementary Information (tableS-1, S-2, S-3, S-4, S-5, S-6). The nucleotide sequences generated in this study have been deposited in GenBank under accession numbers PX763550–PX763551.

## Acknowledgements

This work has been supported by National Natural Science Foundation of China (32372516, 32230086) and the Key Projects of Ningbo Natural Science Foundation (2023j015).

## Author contributions

Ji-Chong Zhuo conceived and designed the study, wrote the manuscript, participated in experiments, and analyzed the data. Hai-Qiang Wang performed most of the experiments and contributed to data analysis. Qing-Ling Hu contributed to the single-cell analysis and assisted with other experiments, along with Ze-Ping Mao, Lin Wang, and Fan Wu. Jia-Bao Lu, Hai-Jian Huang, Zong-Tao Sun, Fei Yan, Jian-Ping Chen, and Jun-Min Li discussed the data and provided partial materials. Chuan-Xi Zhang supervised, organized, and directed the project.

## Extended data

**Extended Data Figure 1.**
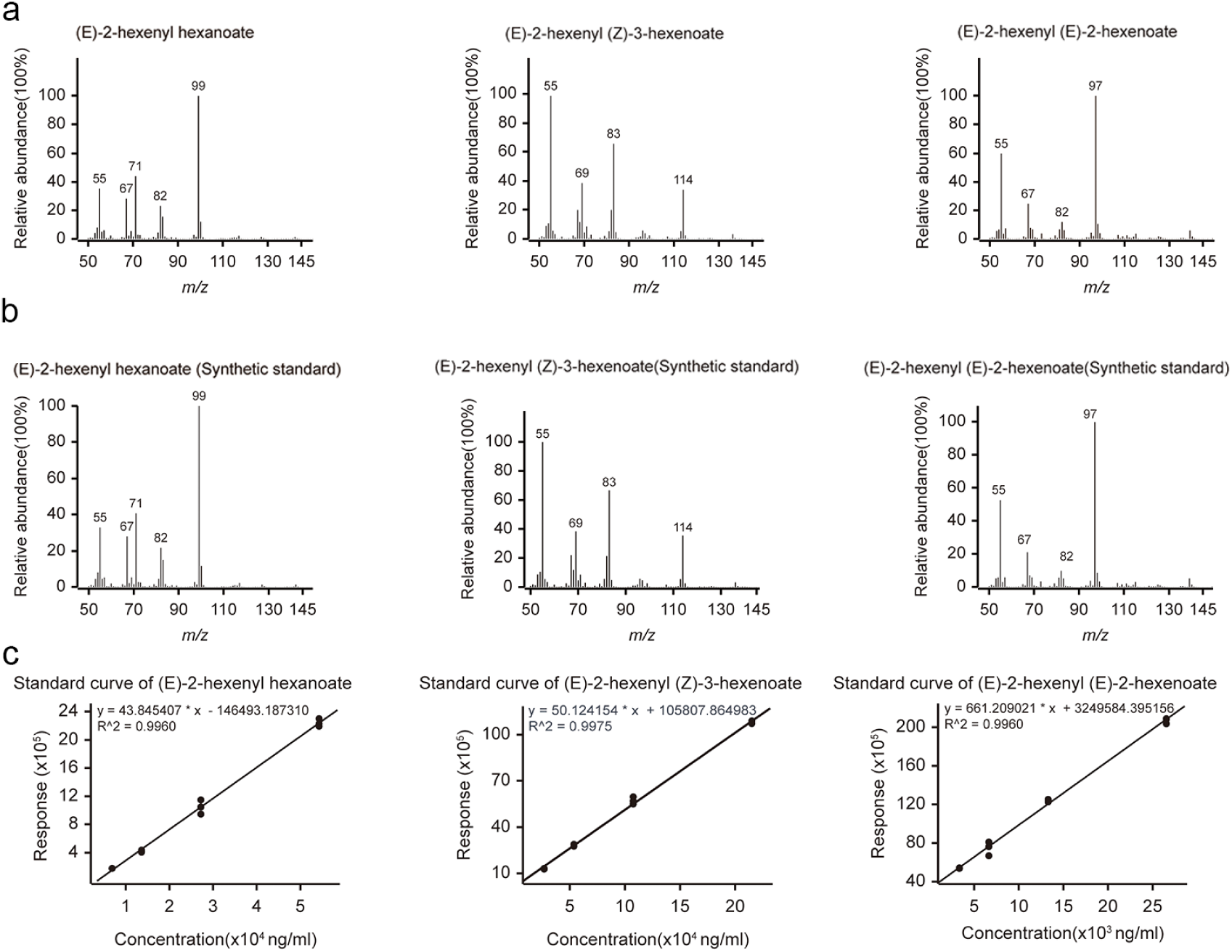
GC–MS identification and quantitative analysis of E2HH, E2HE2H, and E2HZ3H in synthetic standards and biological samples. a,b) GC–MS profiles of synthetic standards and biological samples showing the characteristic peaks of E2HH, E2HE2H, and E2HZ3H. Compounds were identified based on retention times and mass spectra by comparison with authentic standards. This analysis confirms the presence or absence of these compounds in the samples and provides reference data for subsequent quantitative measurements of pheromone components in both females and males. c) Calibration curves of E2HH, E2HZ3H, and E2HE2H for quantitative analysis. Standard curves were generated using synthetic standards at a range of concentrations by plotting peak area against concentration. The resulting regression equations (R² > 0.99) were used for quantitative determination of target compounds in GC–MS analyses of biological samples.

**Extended Data Figure 2.**
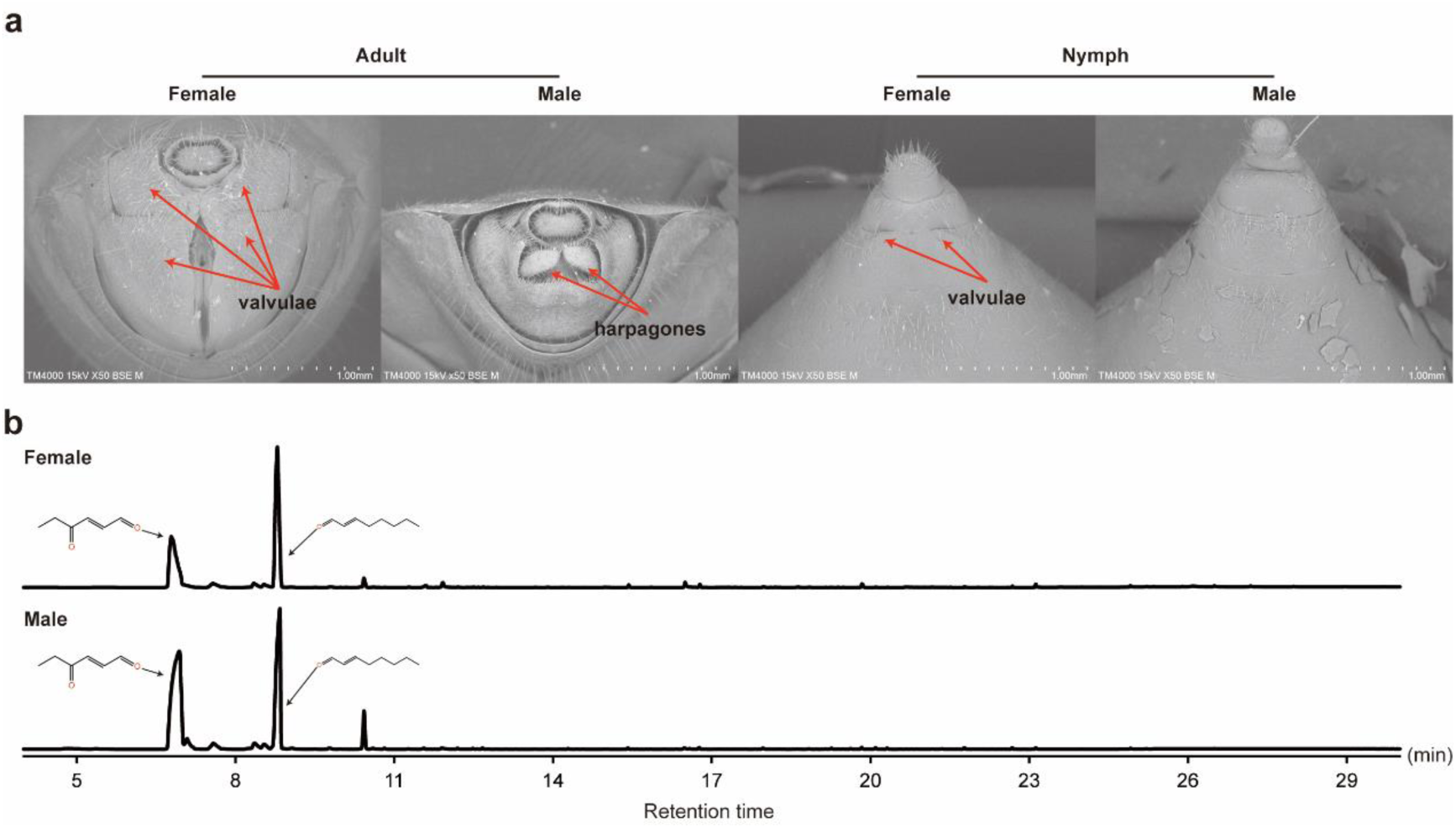
Terminalia of adults and fifth-instar nymphs and GC–MS analysis of fifth-instar nymphs. a) The obvious genital segments differences between of females and males of adult stage or 5^th^-instar nymphs. The female’s valvulae in adult stage and 5^th^-instar nymphs, the male’s harpagones in adult stage are annotated. b) Absence of E2HH, E2HE2H, and E2HZ3H in the GC–MS profiles of fifth-Instar female and male nymphs.

**Extended Data Figure 3.**
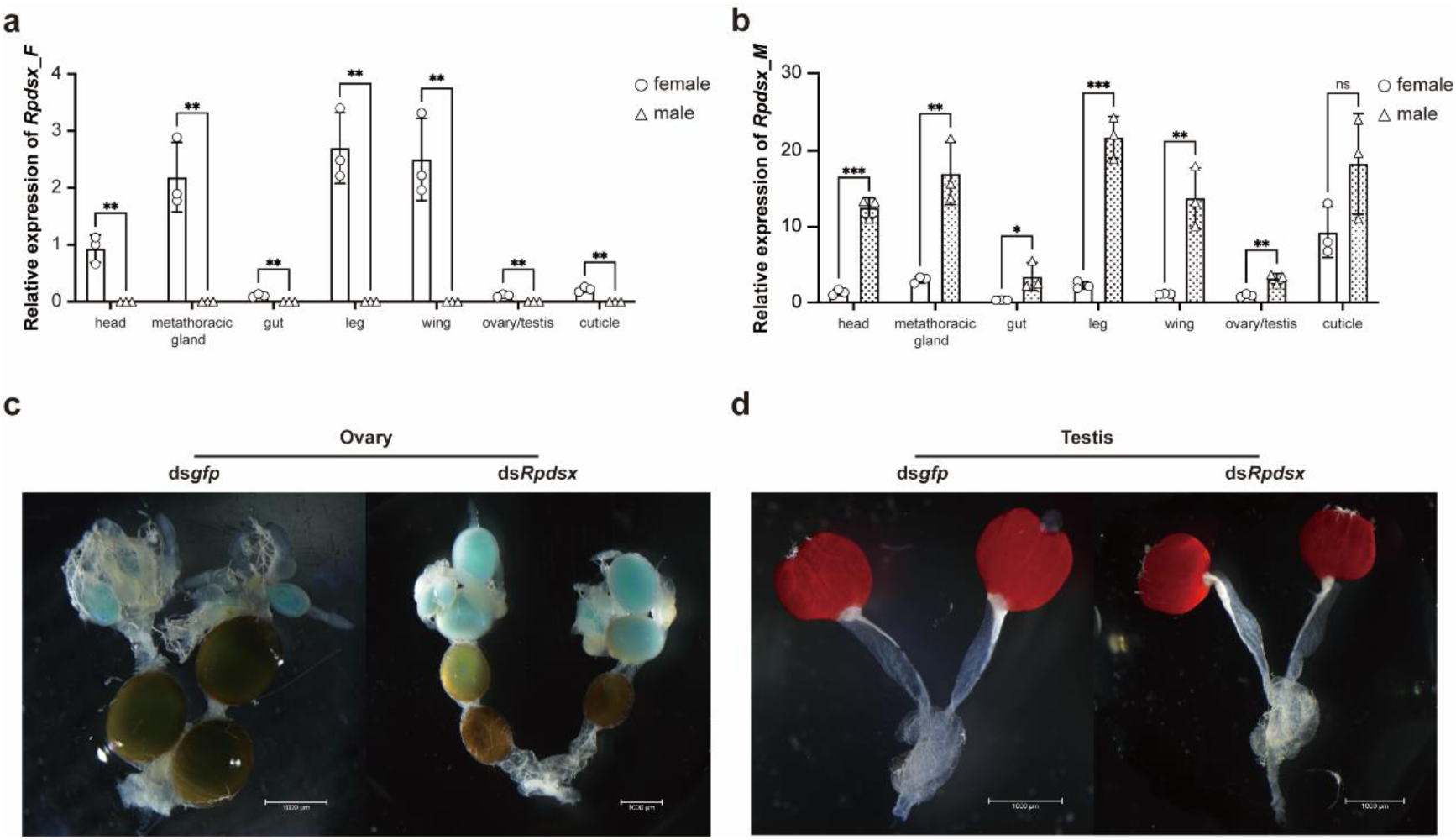
The expression and influence of *Rpdsx* isoforms. a, b) Expression patterns of *Rpdsx_F* and *Rpdsx_M* in the head, metathoracic glands, gut, legs, wings, ovaries/testes, and cuticle of adults. Both *Rpdsx_F* and *Rpdsx_M* exhibited sex-specific expression in the metathoracic glands. c, d) Effects of *Rpdsx* on gonadal development in *R. pedestris*. ds*Rpdsx* or ds*gfp* was injected into newly emerged 1-day-old females and males. After 7 days, ovaries and testes were examined, and no obvious differences were observed between ds*gfp*-treated and ds*Rpdsx*-treated ovaries or testes, respectively.

**Extended Data Figure 4.**
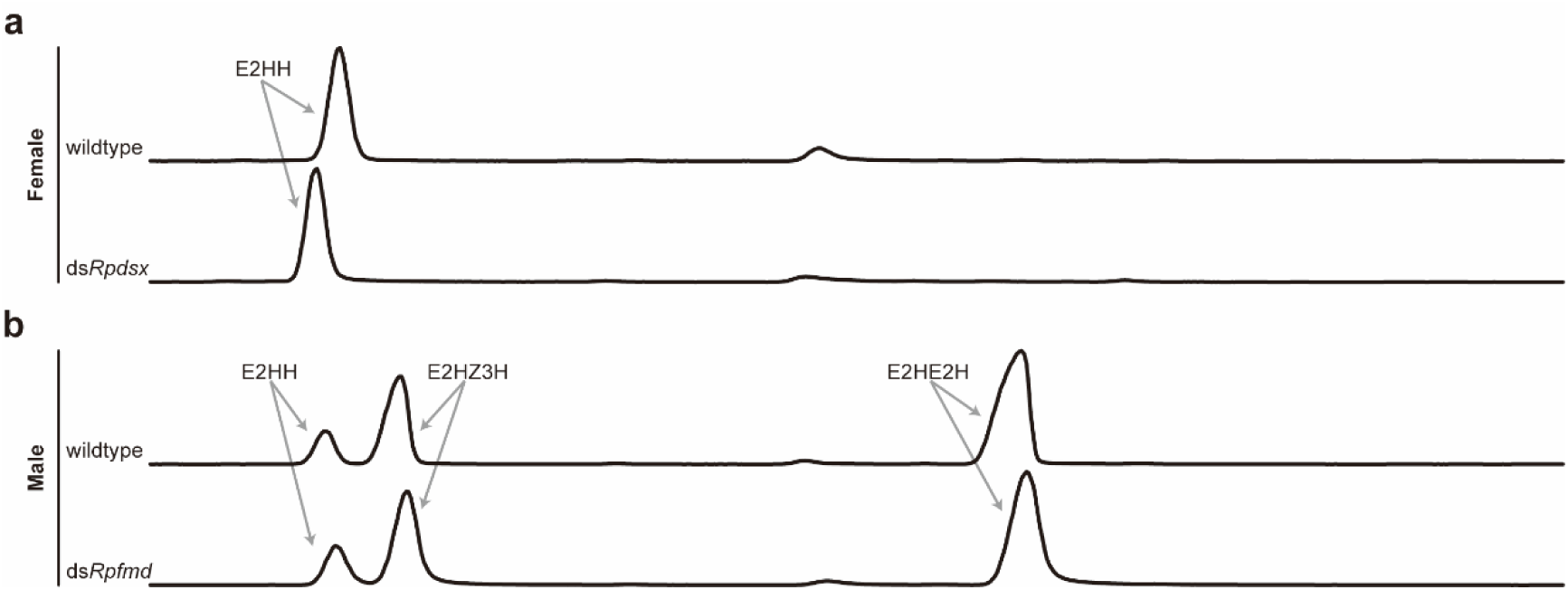
Effects of *Rpdsx* and *Rpfmd* on pheromone production. a) GC–MS profiles of wild-type and ds*Rpdsx*-treated females. Newly emerged 1-day-old females were injected with ds*Rpdsx*, and pheromone levels were analyzed 7 days later. No significant differences were observed compared with wild-type females. b) GC–MS profiles of wild-type and ds*Rpfmd*-treated males. Newly emerged 1-day-old males were injected with ds*Rpfmd*, and pheromone levels were analyzed 7 days later. No significant differences were observed compared with wild-type males.

**Extended Data Figure 5.**
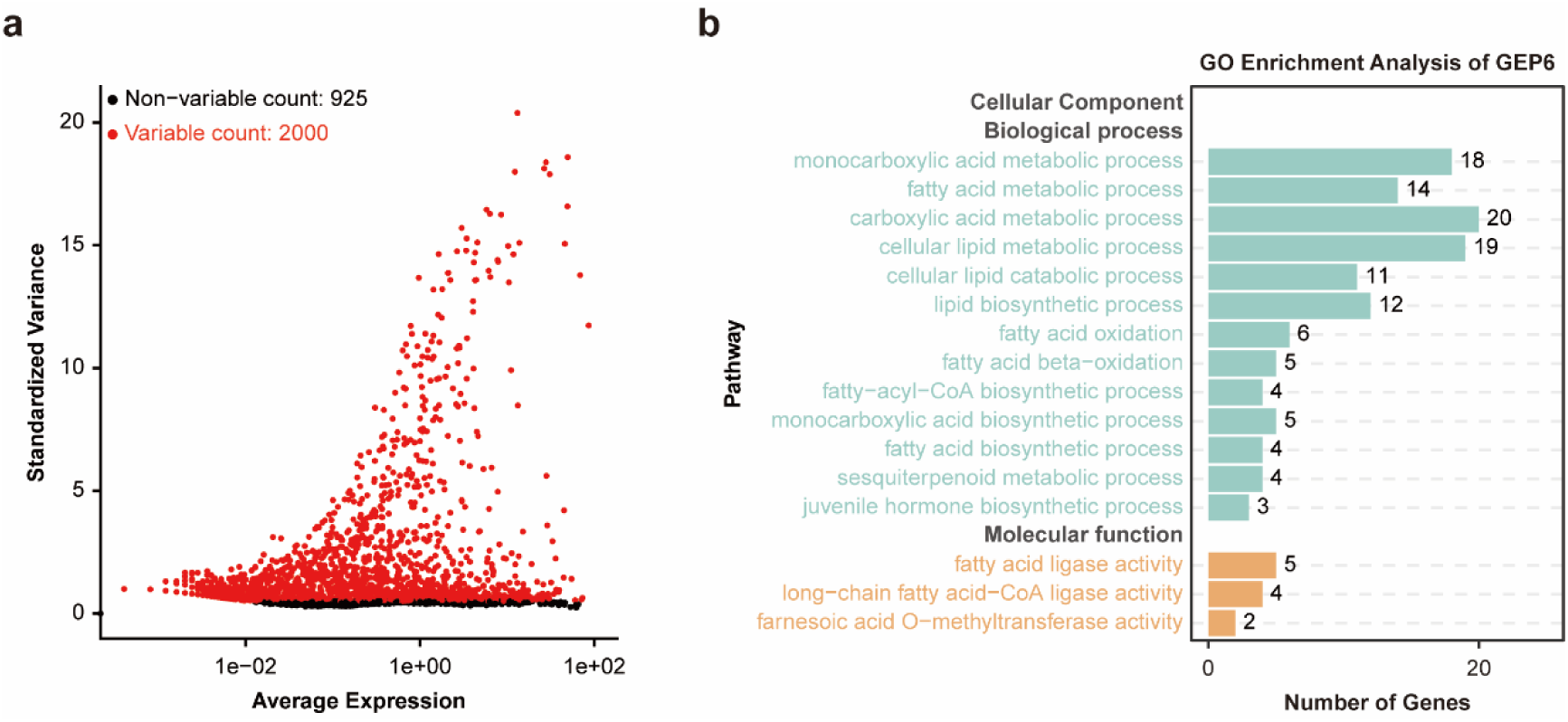
Single-cell analysis. (a) Variable feature plot displaying the 2,000 most variable transcripts selected for all downstream analyses and clustering. These features were used to identify transcriptional heterogeneity across the single-cell dataset. (b) Gene Ontology (GO) enrichment analysis of GEP6, highlighting the overrepresented biological processes, molecular functions, and cellular components in cell type 9.

**Extended Data Figure 6.**
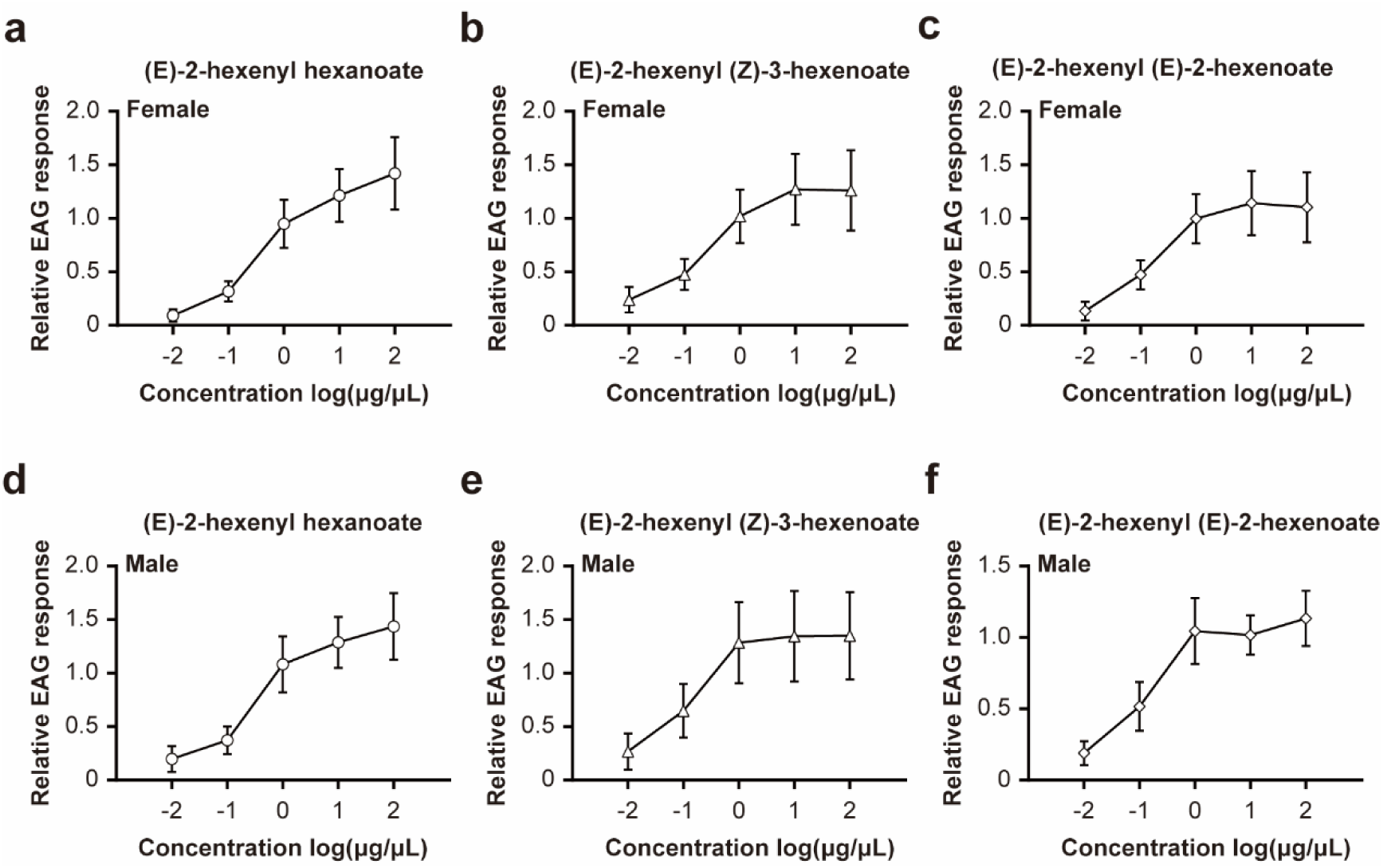
Detection of E2HH, E2HZ3H, and E2HE2H at different concentrations in female (a-c) and male insects (d-f). Synthetic standards of E2HH, E2HZ3H, and E2HE2H were prepared at varying concentrations and presented to adult females and males. This analysis provides insight into the dose-dependent detection of these pheromone components in both female and male insects.

**Extended Data Figure 7.**
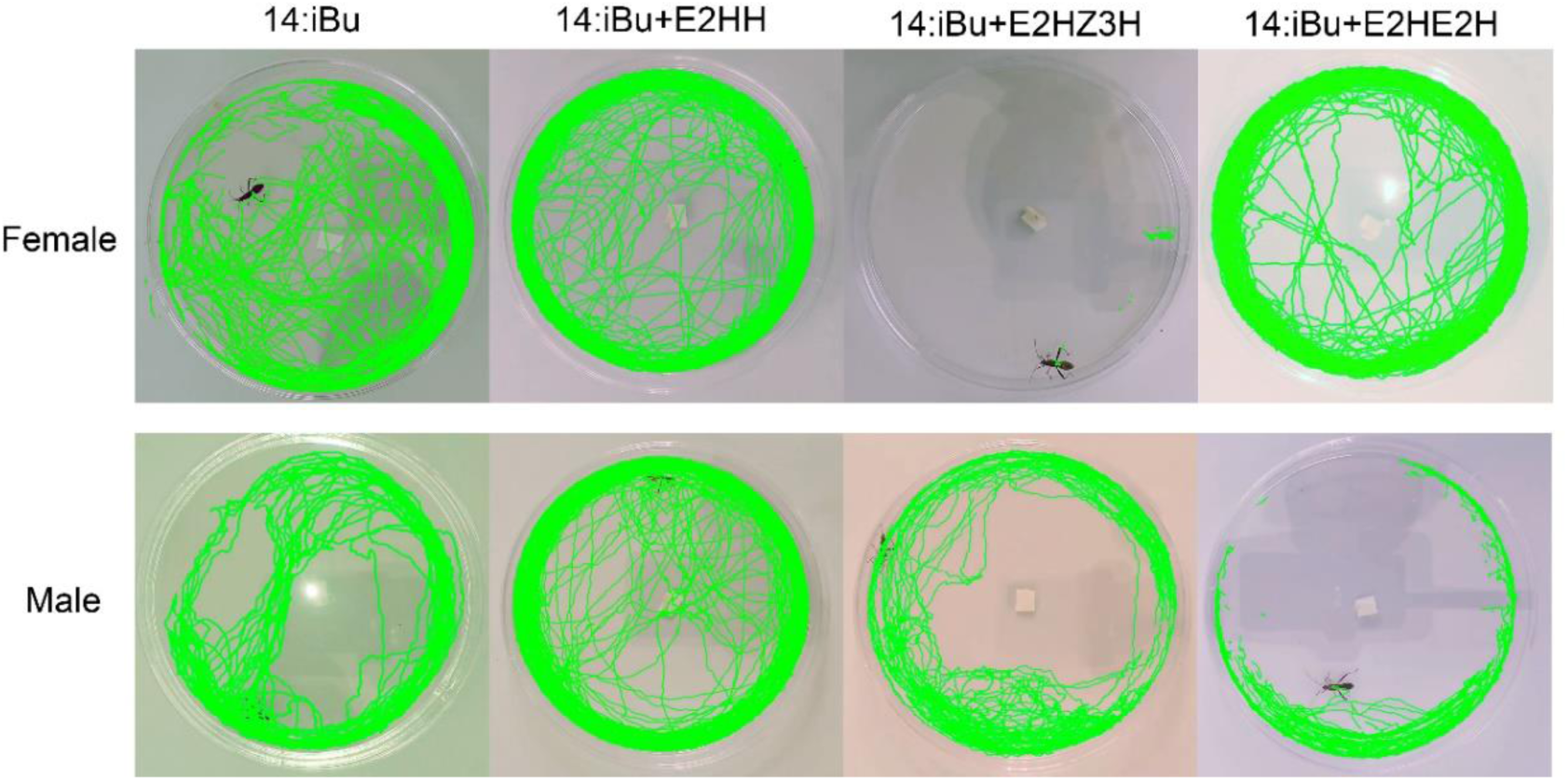
The effects of E2HH, E2HZ3H, and E2HE2H on wild-type females and males were evaluated in 9 × 1.5 cm culture dishes. The green lines represent the movement trajectories recorded over approximately 180 minutes. A 0.5 × 0.5 cm piece of filter paper loaded with 10 μL of 10% 14:iBu, 14:iBu + E2HH, 14:iBu + E2HZ3H, or 14:iBu + E2HE2H was placed at the center of each dish.

Supplementary **Video1. The mating behavior of wildtype males with treated females.** (A) Mating behavior between wildtype female and wildtype male. (B) Mating behavior between wildtype female treated with 1:9 (v/v) E2HZ3H/n-hexane and wildtype male. (C) Mating behavior between wildtype female treated with 1:9 (v/v) E2HE2H/n-hexane and wildtype male. (D) Mating behavior between wildtype female treated with 5% E2HZ3H+5% E2HE2H and wildtype male. The mating happens in 9X1.5cm culture dishes.

Supplementary **Video 2. Mating behavior between wildtype males with treated males.** (A, B) The influence of E2HZ3H or E2HE2H on the mating behavior between wildtype males and ds*Rpdsx*-treated males. (a1, b1,) Wildtype males shows mating behavior to ds*Rpdsx*-treated mlaes. (a2, b2) Adding 10% E2HZ3H or E2HE2H on the body of ds*Rpdsx*-treated mlaes could disturb the recognition of wildtype males, and no mating behavior could be found in 3 hours. (a3, b3) Wildtype males exhibit mating behavior toward ds*Rpdsx*-treated males again after 24 hours, when E2HZ3H or E2HE2H volatilization ceases. The same pair of wild-type and ds*Rpdsx*-treated males (labled by “1”) was used in experiments A and B, respectively. Treated males were marked with a white marker respectively. The mating behaviors are detected in 3.5X1.5cm culture dishes.

